# polyCluster: Defining Communities of Reconciled Cancer Subtypes with Biological and Prognostic Significance

**DOI:** 10.1101/228551

**Authors:** Katherine Eason, Gift Nyamundanda, Anguraj Sadanandam

**Author notes:** To whom correspondence should be addressed, Anguraj Sadanandam, Ph.D., Institute of Cancer Research (ICR), 15 Cotswold Road, Sutton, Surrey, SM2 5NG, United Kingdom.

## Abstract

To stratify cancer patients for most beneficial therapies, it is a priority to define robust molecular subtypes using clustering methods and “big data”. If each of these methods produces different numbers of clusters for the same data, it is difficult to achieve an optimal solution. Here, we introduce “polyCluster”, a tool that reconciles clusters identified by different methods into context-specific subtype “communities” using a hypergeometric test or a measure of relative proportion of common samples. The polycluster was tested using a breast cancer dataset, and latter using uveal melanoma datasets to identify novel subtype communities with significant metastasis-free prognostic differences. Available at: https://github.com/syspremed/polyClustR

## Background

Recently, advances in omics technologies have lead to large volumes of data being collected on molecular profiles, including gene expression, in various cancers. Cancers of all types exhibit inter-tumoral (between patient) heterogeneity that can be quantified in part by gene expression. This heterogeneity can help explain the differential prognosis in cancer patients treated with the same therapies. A well-established example is the specific efficacy of trastuzumab (Herceptin) in HER2-positive breast cancer [1]. Previously, we have suggested potential differential cetuximab (anti-EGFR therapy) response in colorectal cancer (CRC) subtypes [2]. More recently, trials of oxaliplatin in Stage II and III CRC found that its effectiveness may be limited to certain subtypes published by us [2, 3]. In pancreatic cancer, we observed a relatively increased response to gemcitabine in quasi-mesenchymal (QM) subtype cell lines compared to classical subtype cell lines [4]. This result corroborates with the finding by Mofitt *et al.*, that patients from basal-like pancreatic cancer subtype (equivalent to our QM subtype) has improved response to adjvant therapy compared to classical subtype pancreatic tumors [5]. Similarly, we showed potential subtype-specific therapies using a panel of breast cancer cell lines and drug response analysis [6]. Nevertheless, for accurate prediction of therapy responses, the challenge lies in defining robust and clinically relevant subtypes.

In breast cancer, where current opinion lies with the existence of 5 intrinsic gene expression subtypes (basal, HER2/ERBB2, luminal A, luminal B, and normal-like), studies have variously reported a number of subtypes ranging between 4 [7] and 10 [8]. While multiple factors are involved in this apparent discrepancy in defining a number of cancer subtypes, the clustering methodologies can also significantly contribute to this difference. There are various clustering algorithms that are regularly employed for this purpose, and each has its own strengths according to the underlying structure of the data it is applied to. As clustering algorithms have a huge range of potential applications, selection of the appropriate algorithm to use in any given situation can be difficult. At the same time, the need for the user to inspect the results of each algorithm over a range of numbers of clusters (*k*) and select the optimal solution are often subjective. This situation has been improved by the adoption of various consensus clustering techniques, which allow for visual and quantitative examination of multiple re-runs of the same algorithm so the effects of random starting points can be taken into consideration.

However, consensus clustering does not diminish the influence the choice of algorithm has on the clustering solution. The application of different consensus clustering algorithms leads to different number of subtypes (number of clusters, *k*), and hence, defining the optimal number of clusters is often challenging. This is due to various factors in the design of the algorithm: whether it is ‘greedy’, that is, if it makes the locally optimal choice at each individual stage at the possible expense of finding a global optimum; whether cluster centroids must be located at data points; and how iterative algorithms evaluate their convergence to a solution are some examples [9]. This makes the use of a single algorithm to cluster gene expression profiles, as is often done in subtyping studies, risky. In addition, the clusters found may well be valid, but information about either larger stratification of the data or small but distinct sub-subtypes of low frequency may be lost [10]. It is for this reason that finding methods of reconciling optimal clustering solutions identified by different algorithms is necessary. Cluster reconciliation not only validates the clusters from different algorithms – it can also reveal in greater detail the structure in the data on the macro and the micro scale, from broad classifications resulting from a handful of important functional groups, to rarer and less well-defined sub-subtypes. It also reveals more about the efficacy of the clustering algorithms themselves [10, 11].

Here, we demonstrate how to identify optimal solutions and define subtype “communities” by reconciling clusters identified from three different consensus clustering methods - hierarchical clustering (HC) [12, 13], k-means (KM) [14], and non-negative matrix factorization (NMF) [15]. The clusters were further reconciled using at least two approaches. The first, a hypergeometric test to determine the probability that two clusters share the same samples by chance, was previously used to successfully reconcile subtypes of CRC found via clustering in two studies which found three and five optimal subtypes, respectively [2, 10, 16]. It was determined via this analysis that the three subtypes could be appropriately divided into the five sub-subtypes. When four further studies into CRC were published, finding between 3 and 6 optimal clusters [17-20], the Jaccard index was applied to help understand the relationships between these solutions and find “consensus molecular subtypes” (CMS) [11]. The second and a new reconciliation measure used here – calculating the relative proportion of samples in a smaller cluster present in a larger one (termed Eason-Sadanandam index) – differs from measures of cluster similarity such as the Jaccard index in order to give sub-subtypes a high score, even if they are much smaller than a larger cluster (see **Methods** section).

All the above reconciliation methods are part of our new framework or package called “polyCluster”. The framework is flexible that other methods can be included any time. Here, we demonstrate how our new framework can be used to identify breast cancer gene expression “subtype communities” and to compare with existing intrinsic subtypes [7]. Moreover, we have applied this to uveal melanoma gene expression profiles to define novel gene expression “subtype communities” with different prognosis and chromosomal aberrations associated with them.

## Results and Discussion

Our reconciliation method (Figure 1) uses a matrix of preprocessed and normalized gene expression (or any other similar data) and performs the following: a) applies different consenusus clustering methods (including NMF, HC and KM) and uses statistical scores (specific to each method described below) for each clustering to determine the optimal number of clusters; and b) reconciles the results from different clustering methods and identifies a consensus solution by creating network of clusters that defines communities of integrated subtypes using methods such as the hypergeometric test and proportion of maximum intersection (PMI). We then identify the optimal “community” with highest average silhouette width [21] and compare this reconciliation to known subtypes, if they exist, for that set of samples. To illustrate this, we used published gene expression profiles from breast cancer and uveal melanoma as examples.

**Figure 1.**
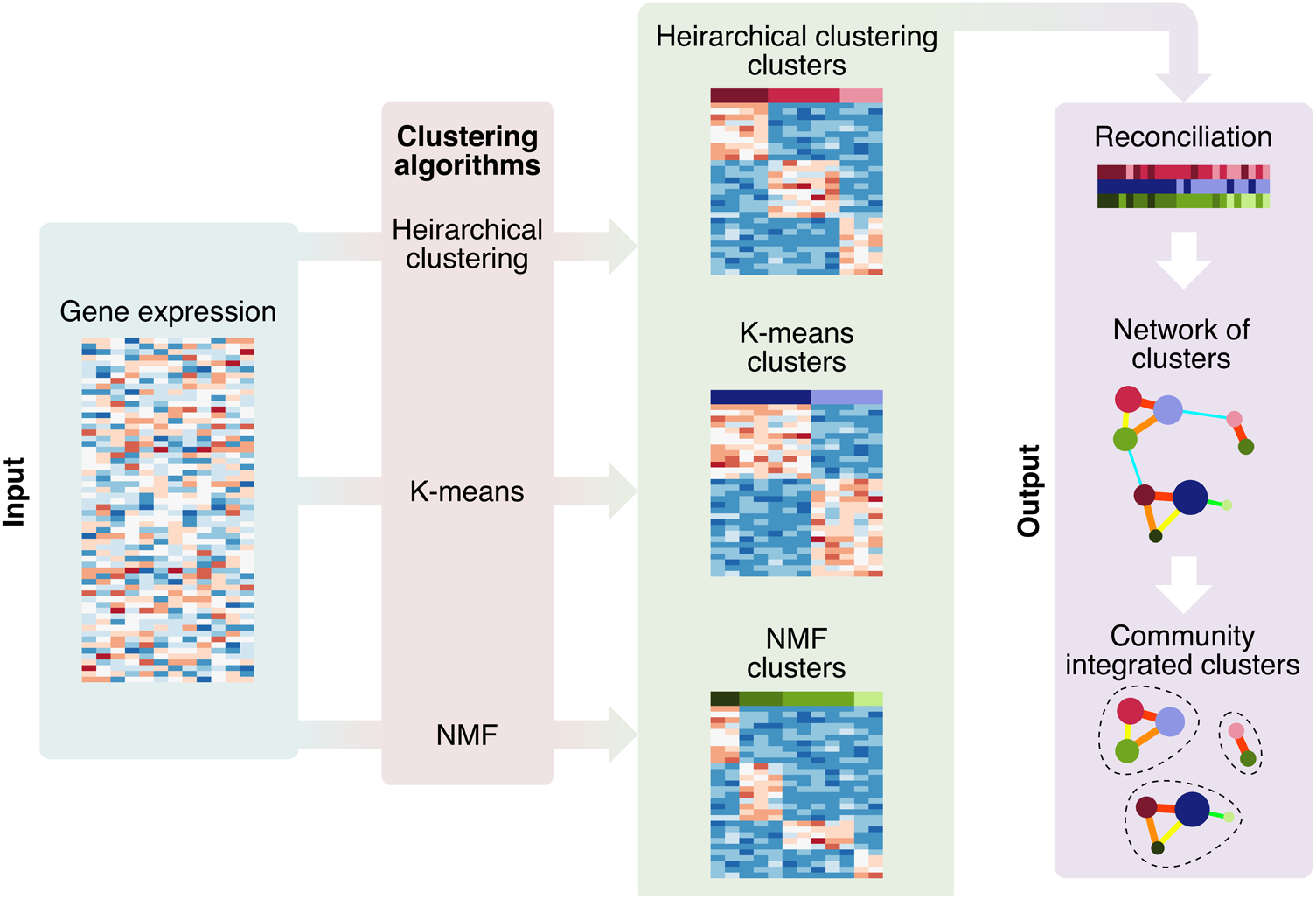
An overview of our pipeline for cluster reconciliation. Gene expression – or other equivalently structured molecular data – is input as a genes by samples matrix. This data is then fed through multiple consensus clustering algorithms (in this case, HC, KM and NMF) to produce multiple clustering solutions. These are then reconciled to create “subtype communities” of similar clusters from across the algorithms’ solutions, by applying community detection to networks representing the similarity between clusters from all the algorithms.

### Application to reconcile breast cancer “subtype communities” with intrinsic subtypes

#### Breast cancer subtypes defined by multiple clustering methods

For this purpose, we used breast tumor gene expression data (*n* = 118) from a published study [22]. Details of initial clustering of this dataset and selection of *k* clusters for each algorithm are provided in **Figures S1A-C**. Initially, we applied the NMF to the 2258 most highly variable genes from this Chin data set as selected by standard deviation (SD>0.8). We identified highest cophenetic correlation coefficient of 0.9997 at *k* subtypes for NMF *k_NMF_*=2 followed by 0.9962 at *k*_NMF_=6. Silhouette width also showed peaks at *k*_NMF_ at 2 and 6 (**Figures S1 A-C**). In order to capture the most heterogeneity, we chose *k*_NMF_=6, and named the clusters breast cancer (b)NMF1 to 6. Overall, known subtypes of these samples [22] were significantly associated with these clusters (Fisher’s exact test; *p*<0.001). Specifically, the clusters bNMF1, bNMF3 and bNMF4 were significantly associated with luminal A, basal and luminal B, respectively (hypergeometric test; false discovery rate; *FDR*<0.01) (Figure 2A). The basal subtype was also border-line significantly associated (*FDR*=0.2) with bNMF5, suggesting the existence of a sub-subtype of basal breast cancer that was not identified earlier when subtypes for this dataset were predicted by correlation with intrinsic subtype signatures [23] [7]. bNMF2 and bNMF6 were not significantly associated with any of the published subtypes. Gene set enrichment analysis (GSEA) of these unidentified subtypes revealed associations with metaplastic breast cancer (bNMF2, *FDR*<0.01) and with 17q21-q25 amplicon gene sets (bNMF6, FDR < 0.1) (**Figure S2A-B**). Overall, application of NMF to the Chin data set identified clusters that partially overlapped with published subtypes, and others with interesting breast cancer biology.

**Figure 2.**
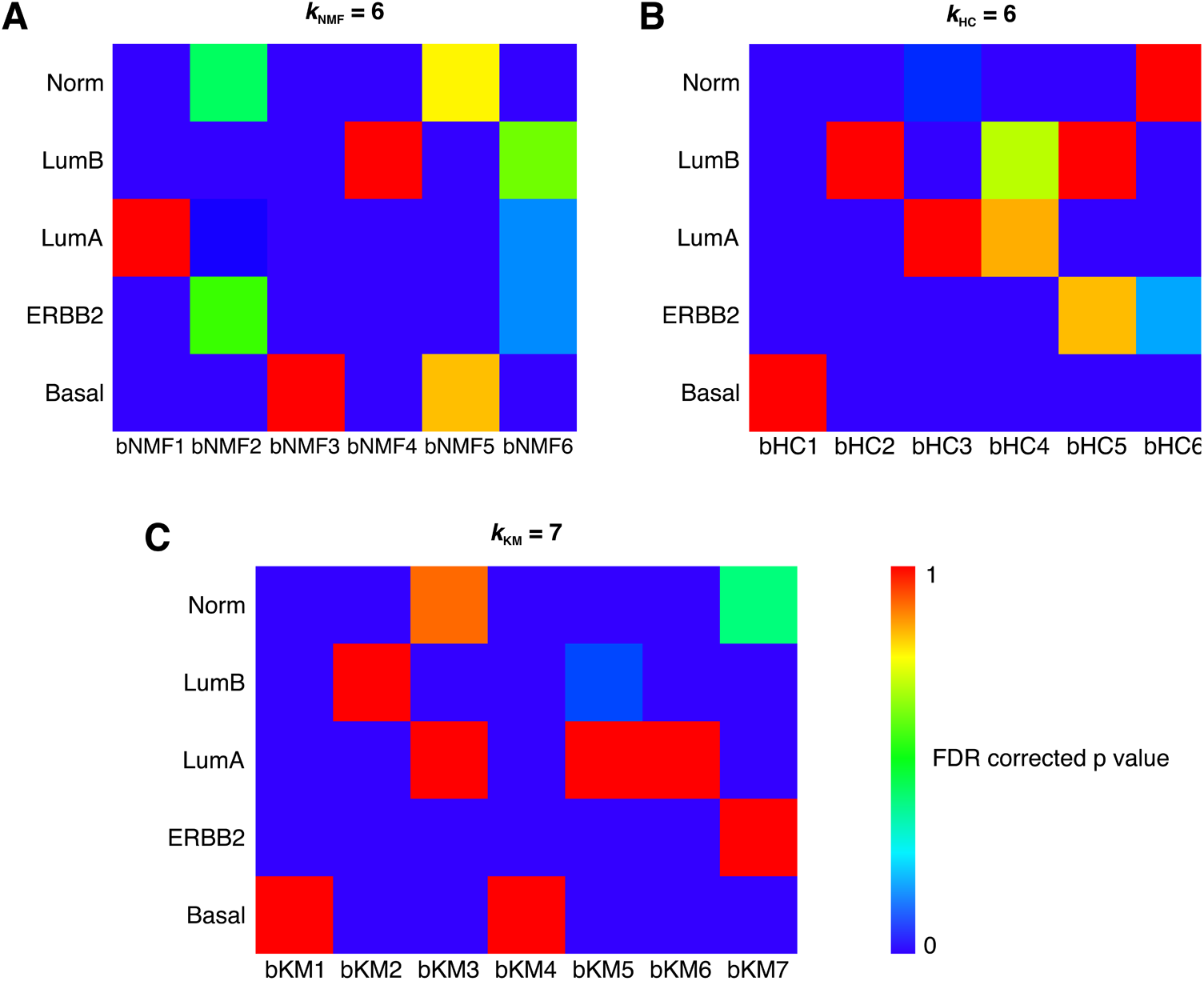
Breast cancer subtypes and their association with intrinsic subtypes – application of polyCluster. (**A-B**) Similarity of each set of clusters generated by consensus A) NMF, B) HC and C) KM to the known breast cancer subtypes of each sample (as assigned by correlation to PAM centroids) using 118 breast cancer samples from a published dataset [22]. A hypergeometric test was used to test the significance of overlap between the clusters and the known subtypes. bNMF, bHC and bKM represent NMF, HC and KM subtypes, respectively. Norm – normal-like, lumA – luminal A and lumB – luminal B subtypes.

Since NMF identified extra subtypes in Chin data set, we applied two additional clustering methods – consensus hierarchical clustering (HC) and K-means (KM). When we applied consensus hierarchical clustering to the same data, *k*_HC_=2 and *k*_HC_=6 had the highest silhouette widths. (**Figures S1A and C**). The cophenetic coefficient after *k*_HC_= 6 does not increase significantly and the consensus plot showed consensus clusters (**Figures S1A and C**). Hence, we chose six HC clusters to again cover the most heterogeneity. The clusters from HC for breast cancer data were defined as breast cancer (b)HC. As with the NMF clusters, these clusters were significantly associated with the known subtypes of these samples (Fisher’s exact test; *FDR*<0.001). The bHC1, bHC3 and bHC6 clusters were significantly (hypergeometric test; *FDR*<0.01) associated with basal, luminal A and normal-like subtypes, respectively (Figure 2B). Both bHC2 and bHC5 were significantly (*FDR*<0.01) associated with luminal B. bHC4 was marginally significantly associated with luminal A subtype, and bHC5 with the ERBB2 (HER2) subtype, with less significance (*FDR*<0.2; Figure 2B).

Additionally, we applied consensus KM clustering to the Chin data set. While both the cophenetic coefficient and silhouette width showed highest peaks at *k*_kM_=3 and 4 (after *k*_KM_=2), we observed that consensus clustering at these *k*_KM_s did not show clear consensus clusters. There were not large differences in cophenetic coefficient, silhouette width and consensus clusters at *k*_KM_ between 4 and 7 (**Figure S1A and D**). Hence, we chose *k*_KM_=7 as an optimal cluster. All of these KM clusters (defined as breast cancer (b)KM were significantly associated with known breast cancer subtypes (Figure 2C; Fisher’s exact test; *p* < 0.001), unlike the NMF and HC clusters. Specifically, bKM1 and bKM4 were associated with basal, bKM2 with luminal B and bKM3, bKM5 and bKM6 with luminal A (hypergeometric test; *FDR* < 0.01). bKM7 was significantly associated with the ERBB2 subtype, which was not highly significant with any NMF or HC clusters. bKM3 was marginally associated with the normal-like subtype (*FDR*=0.08). Direct comparison of the two basal clusters through GSEA revealed enrichment of multiple gene sets associated with invasive breast cancer, immunity and cytokines (**Figure S2C-F**). This clearly suggests that different clustering algorithms have the inherent capacity to identify distinct clusters. Here, KM has identified clusters with more significant association to published subtypes.

#### Identification of breast cancer “subtype communities”

The existence of multiple clustering solutions defined by different algorithms poses the question of what number of clusters is optimal, and how they reconcile between different methods. To address these questions, we chose two different reconciliation methods – hypergeometric test and proportion of maximum intersection. The results from each of the reconciliation methods are discussed below.

Previously, we have used the hypergeometric test to assess enrichment of samples between two CRC classifications (including ours) as a means of reconciling subtypes [10]. Similarly, we have used this analysis here to reconcile breast cancer clusters between the three different (NMF, HC and KM) algorithms utilized above. Subsequently, in order to group those clusters with significant similarity into “subtype communities”, we performed network community detection by applying weighted label propagation method (using FDR values as edge weights) [24]. As a result, we observed six “subtype communities” (groups of clusters; bHYP1-6) based on this analysis (Figure 3A).

**Figure 3.**
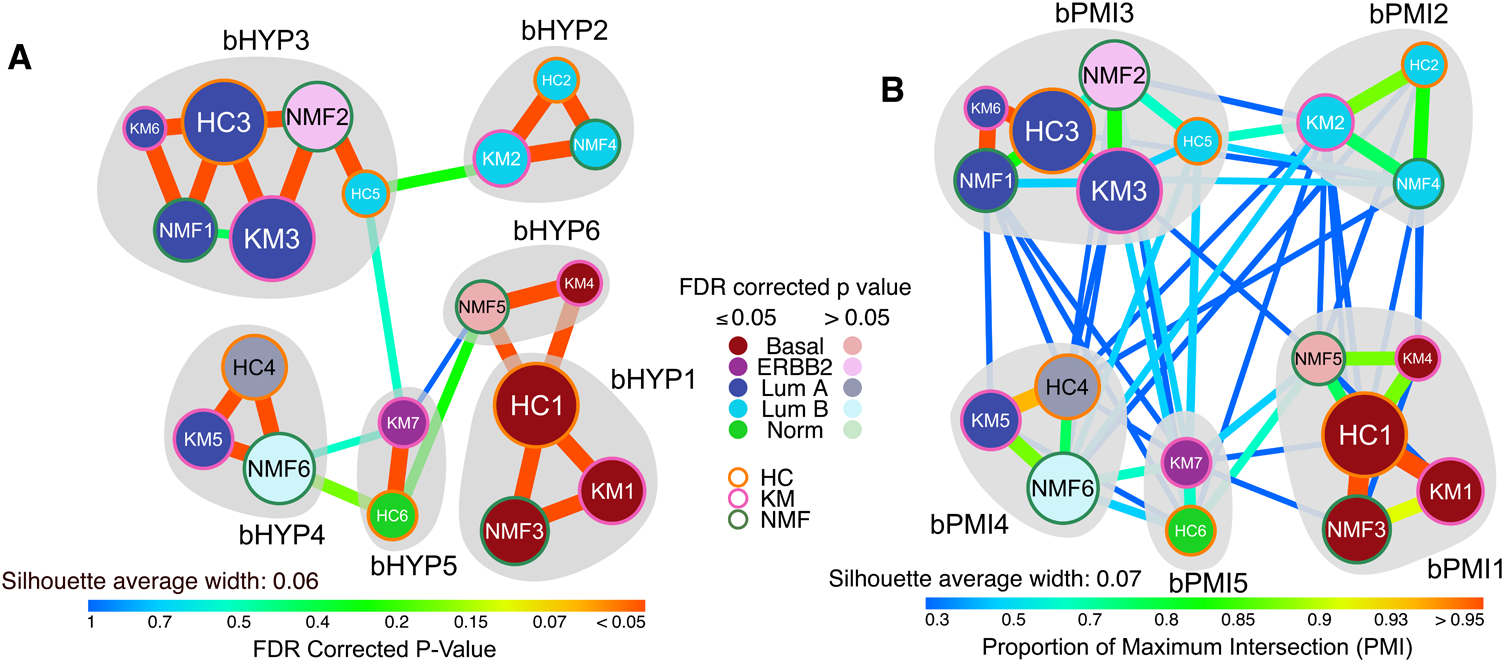
Subtype communities of breast cancer identified using polyCluster. (A) Ahypergeometric (HYP) test and (B) PMI was used to assess the significance of the overlap between each pair of clusters using Chin breast cancer data set. The resulting FDR corrected p values were plotted as edge colours/weights in this network, with each node representing a cluster. The size of each node represents the number of samples that cluster contains, and those nodes in a lighter shade represent clusters with associations to known subtypes that are not significant (FDR corrected *p* > 0.05). Gray shading marks dense groups of clusters as defined by network community detection. bHYP and bPMI represent HYP and PMI subtype breast cancer communities, respectively.

There was significant association with the known subtypes and these communities (Fisher’s exact test; *p*<0.001). We observed that five communities were primarily and significantly (hypergeometric test; *FDR*< 0.05) associated with published breast cancer subtypes – bHYP3 and bHYP4 with luminal A, bBHYP2 with luminal B and bHYP1 and bHYP6 with basal (Figures 3A and S3A). Four of the communities (bHYP1-4) contained clusters from all three clustering algorithms (Figure 3A). Interestingly, each of the luminal A and basal subtypes were split into two communities. One basal community (bHYP6) contained the immune-enriched bKM4 cluster. One of the luminal A communities (bHYP3) contained a number of samples from the ERBB2 subtype in a cluster that was enriched for a metaplastic breast cancer signature (bNMF2; Figures 3A and S3A), while the other (bHYP4) contained some luminal B samples in the 17q21-q25 amplicon-enriched cluster (bNMF6; Figures 3A and S3A). Finally, there was a community (bHYP5) with mixture of normal-like and ERBB2 subtype samples. This community was the most mixed in terms of intrinsic subtypes. Overall, hypergeometric test-based reconciliation expanded the breast cancer subtypes to 6 communities.

Our PMI method is similar to the Jaccard analysis that we used recently to reconcile CRC subtypes as a part of the CRC Subtyping Consortium (CRCSC) [11], with the difference that it weights sub-groups of a larger cluster as strongly as identical clusters of the same size (see **Methods**). Here, we applied the PMI method to reconcile subtypes from NMF, HC and KM similar to what we performed using the hypergeometric test. Unlike the hypergeometric method, PMI identified five communities (bPMI1 to 5; Figures 3B and **S3B**), four (bPMI2 to 5) of which were analogous to hypergeometric communities (bHYP2, 3, 4 and 5). The final community (bPMI1) was a combination of the two basal hypergeometric communities (bHYP1 and 6). These communities were significantly associated with known subtypes, overall (Fisher’s exact test; *p* < 0.001). As expected, four of the five communities represent luminal A (bPMI3 and 4), luminal B (bPMI2) and basal (bPMI1) communities (hypergeometric; *FDR*<0.05). The other community (bPMI5) was a mixture of HER2/ERBB2 and normal-like (Figures 3B and **S3C**).

To chose optimal “subtype community” between HYP and PMI communities, we calculated the silhouette width [21] for all samples in the different communities (Figures 3 and **S4**). The average silhouette widths for HYP communities were 0.06 and that for PMI communities were 0.07. Hence, PMI communities with highest average silhouette width were chosen as optimal.

This application of the pipeline to a well-characterised cancer has demonstrated its ability to identify new biologically distinct “subtype communities” of patients, alongside those subtypes which have already been extensively described. We next sought to apply this pipeline to a cancer with molecular subtypes that have not been explored so comprehensively, although uveal melanoma classes at gene expression levels are known [25-27].

#### Application to uveal melanoma and identification of novel “subtype communities”

##### Identification of subtype communities

Compared to breast cancer, uveal melanoma is a cancer type that has not been extensively subtyped, presumably due to its low incidence. This scarcity of samples makes clustering a challenge – clusters discovered are less likely to be robust due to their small size. It is in cases such as this where the reconciliation of clusters from multiple algorithms may present benefits in terms of increasing confidence in the results of clustering.

As with the breast cancer data, we applied the three clustering algorithms of HC, KM and NMF to a dataset of the 6146 most variable genes (SD>0.8) from 58 patients with uveal melanoma (GSE22138, [28]). By performing the same assessment of cophenetic coefficient, silhouette width and consensus matrices, we discovered four clusters by HC, six clusters by KM and five clusters by NMF (**Figure S5A-D**). This demonstrates that different clustering methods yield different clusters using the same data set. However, reconciling the results from these methods to identify the optimal number of clusters can characterize the more heterogeneity in uveal melanoma that may be associated with disease phenotypes such as metastasis and abnormalities in chromosome 3.

By reconciling these subtypes by a hypergeometric test followed by community detection, we identified five “subtype communities” of clusters (Figure 4A). When we assessed these communities for the key molecular feature of chromosome 3 aneuploidy, we discovered a significant association of these communities with this feature (Fisher’s exact test; *p*<0.001); one community – melanoma mHYP2 – was significantly enriched (hypergeometric test; *FDR*<0.001) for monosomy, and another (mHYP5) was significantly enriched (*FDR*< 0.05) for both disomy and partial monosomy (Figures 4A and **S6A**). Two of the remaining three communities showed less significant associations with chromosome 3 disomy (mHYP4) and monosomy (mHYP1; hypergeometric test; *FDR*<0.2) respectively, while the final community (mHYP3) was not significantly enriched for either. A similar pattern of associations was observed when assessing four “subtype communities” defined by the PMI method (Figure 4B), with one community each representing monosomy and disomy (mPMI1 and mPMI4, respectively), and one mixed disomy/partial monosomy/monosomy community (mPMI2) – however the association was not statistically significant (Fisher’s exact test; *p*=0.577). (Figures 4B and **S6B**). HYP subtypes were chosen over PMI subtypes for significant association with known key molecular features of uveal melanoma and having lower number of samples with negative silhouette width in this cohort (**Figure S7**).

**Figure 4.**
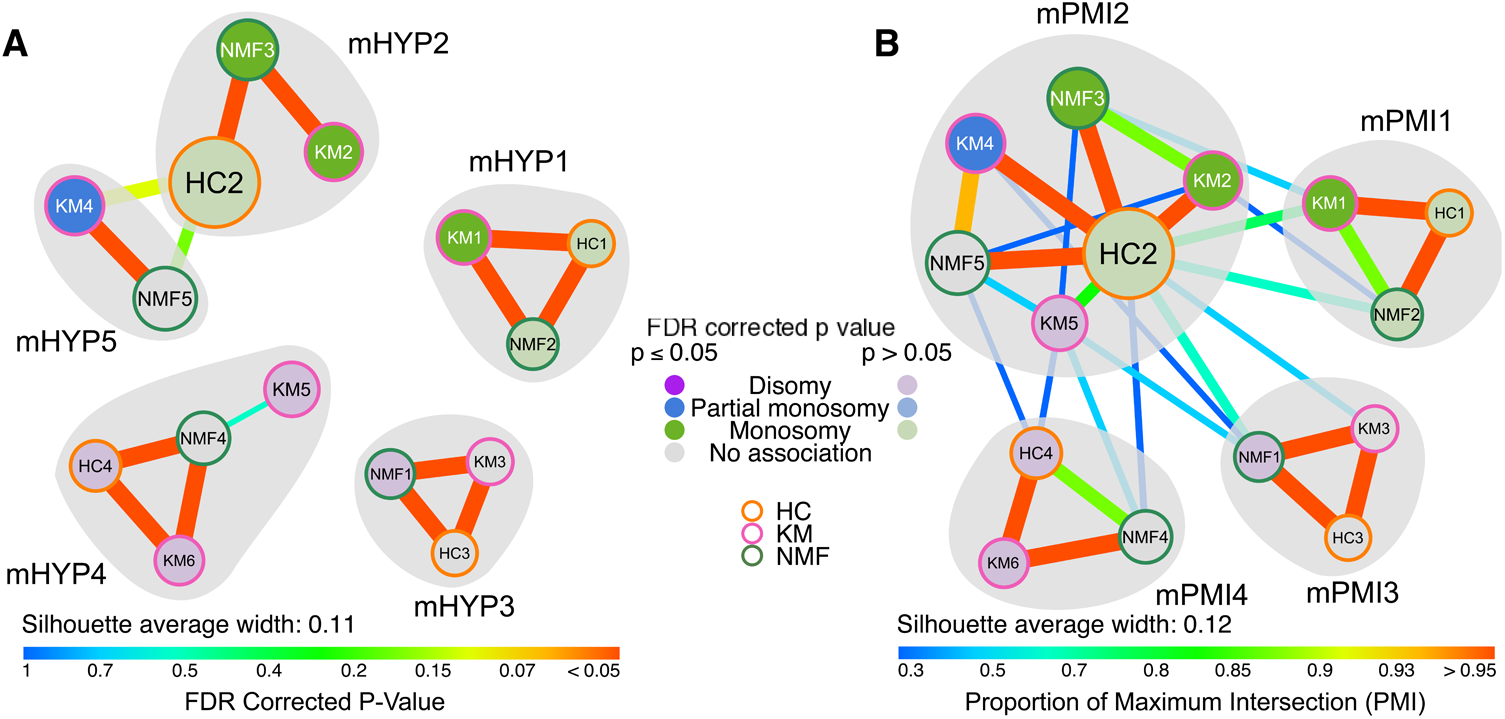
Subtype communities of uveal melanoma identified using polyCluster. (A) A hypergeometric (HYP) test and (**B**) PMI was used to assess the significance of the overlap between each pair of clusters using uveal melanoma data set. The resulting FDR corrected p values were plotted as edge colours/weights in this network, with each node representing a cluster. The size of each node represents the number of samples that cluster contains, and those nodes in a lighter shade represent clusters with associations to known subtypes that are not significant (FDR corrected *p* > 0.05). Gray shading marks dense groups of clusters as defined by network community detection. mHYP and mPMI represent HYP and PMI subtype melanoma communities, respectively.

##### Biological understanding of uveal melanoma subtype communities

Next, we sought to understand these communities by performing GSEA, and discovered that one of these communities (mHYP1) was significantly enriched (*FDR*<0.05) for gene sets associated with immune pathways (e.g. cytokine-cytokine receptor interactions, T cell receptor signaling and JAK-STAT pathway; *FDR*<0.05; Figure 5A–D). On the other hand, another subtype (mHYP3) was associated with neural cell types (e.g. neuron markers, neurotransmitter signaling, neural subtype glioblastoma; Figure 5E–H; *FDR*<0.05). The last communities (mHYP2, mHYP4 and mHYP5) did not significantly associate with any gene sets. This could indicate that mHYP2 enriched for chromosome 3 monosomy and mHYP4 may be by disomy, may be defined by that particular phenotype as opposed to a coherent transcriptomic pattern.

**Figure 5.**
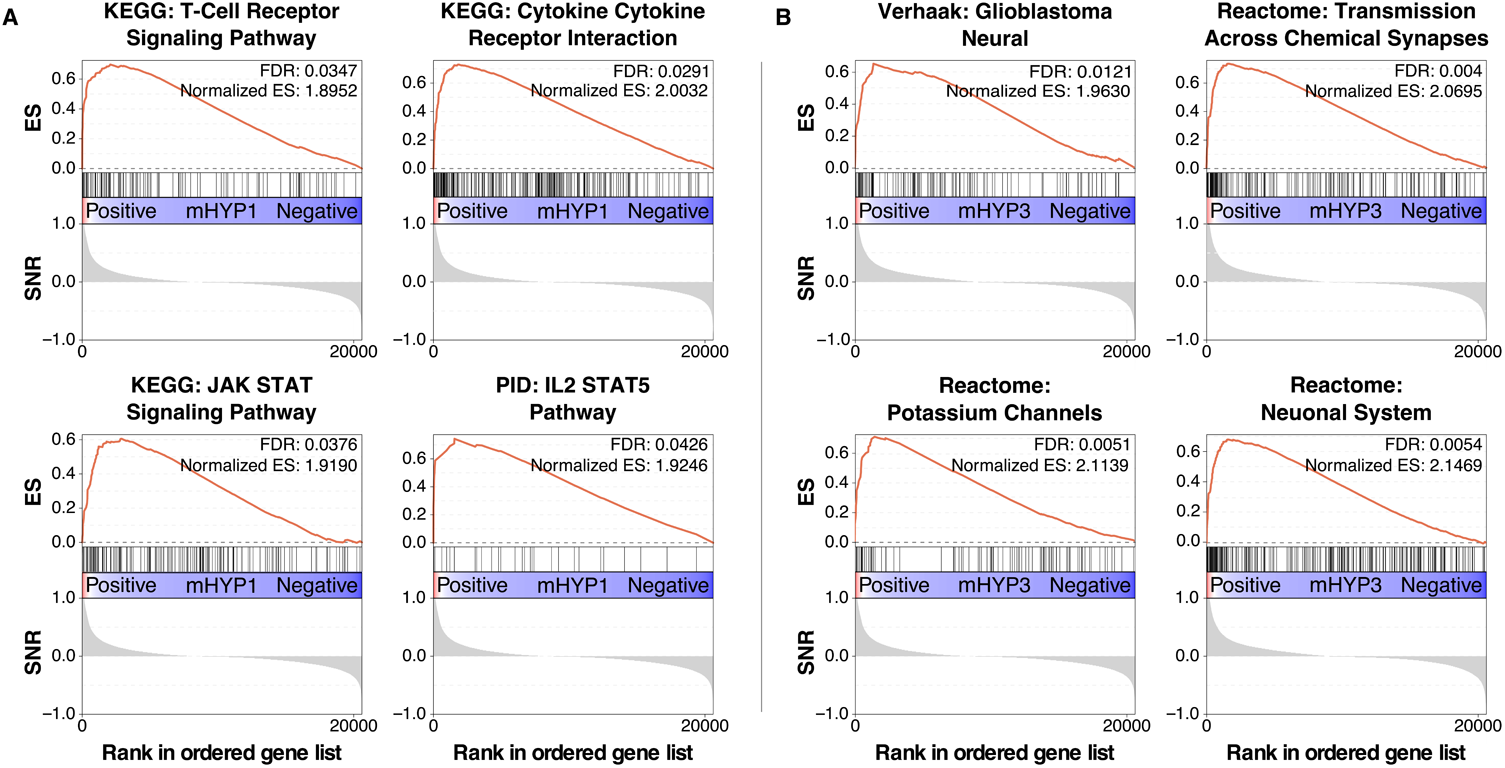
**GSEA enrichment plots** of (A) the mHYP1 uveal melanoma community, showing significant enrichment of immunity-related gene sets, and (B) the mHYP3 uveal melanoma community, showing significant enrichment of neural-related gene sets.

##### Patient prognostic differences between uveal melanoma subtype communities

Since more than 50% uveal melanoma patients undergo metastasis [28], we assessed the metastasis-free prognosis of the uveal melanoma subtype communities using the GSE22138 [28] data set. Among the two highly frequent communities, mHYP2 (37%) showed significantly poorest metastasis-free prognosis, whereas mHYP5 (28%) showed better prognosis. Both mHYP4 (20%) and mHYP1 (11%) communities showed intermediate prognosis (Figure 6A).

**Figure 6.**
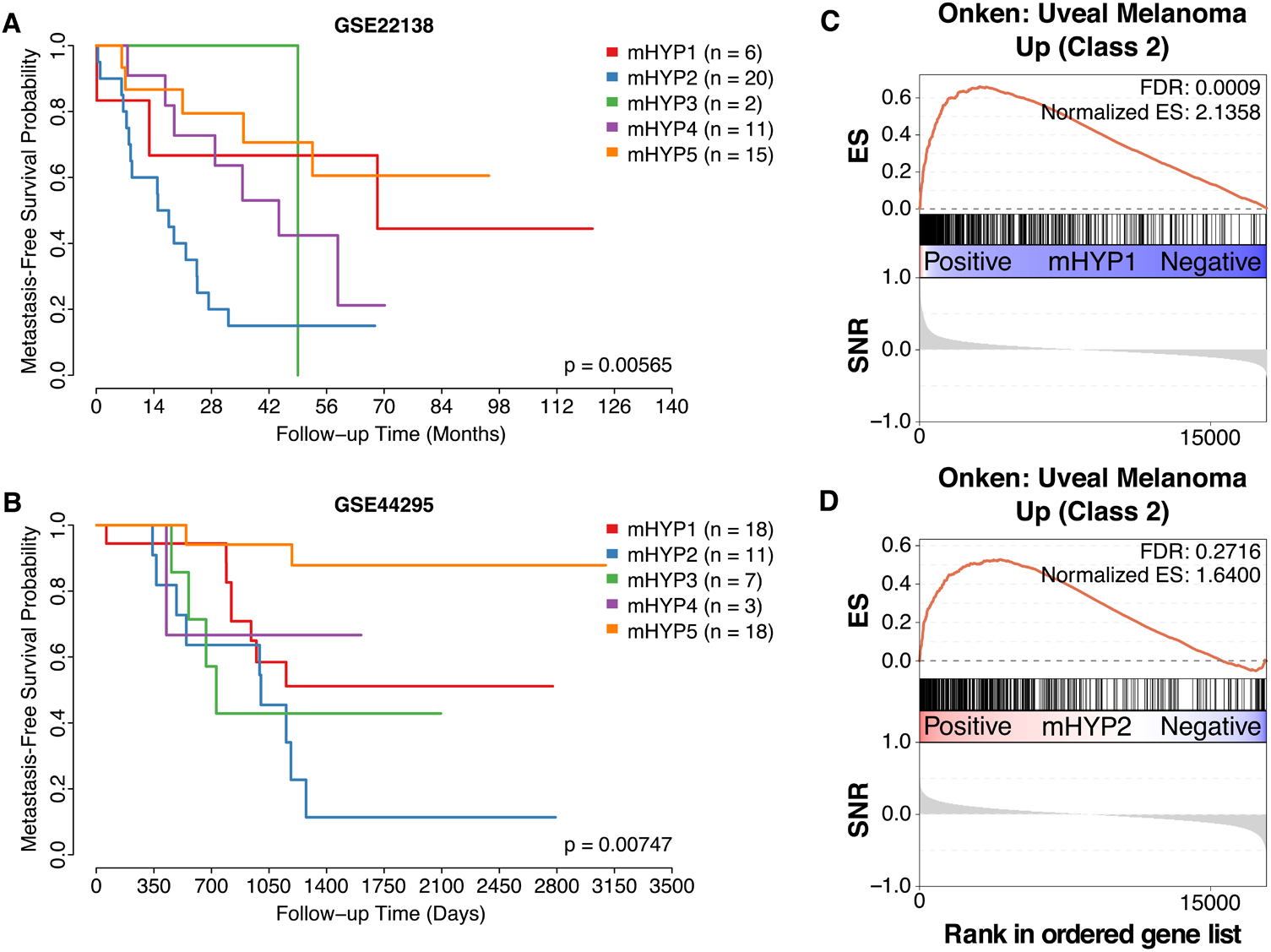
Prognosis and GSEA analysis of uveal melanoma subtype communities. (A-B)Metasisis-free survival in the A) discovery and B) validation cohorts, respectively, was significantly different between communities. (C-D) GSEA enrichment plots of C) the mHYP1 and (B) the HYP3 uveal melanoma communities, showing significant enrichment of class 2 published subtypes,.

##### Validation of uveal melanoma subtype communities

Due to the low frequency of some of these communities in this dataset (5% mHYP3, 11% mHYP1), we sought to validate them in an independent dataset consisting of 58 patients with uveal melanoma (GSE44295). Patients were assigned to subtypes based on the correlation of their gene expression profile with the prediction analysis of microarrays (PAM) [29] centroids of each community. In the validation cohort, 31% of patients were assigned to the mHYP1 (immune-enriched) group, 19% mHYP2 (monosomy-enriched), 14% mHYP3 (neural-enriched), 5% mHYP4 (undetermined) and 31% mHYP5 (disomy/partial monosomy-enriched). In terms of prognosis, these groups showed statistically significant differential metastasis-free survival (*p* = 0.00747; Figure 6B). Analogous to the previous dataset, mHYP2 and mHYP5 communities showed poor and good prognosis respectively. While mHYP1 showed intermediate prognosis, mHYP4 couldn’t be assessed due to low sample size of only 5% (n=3). Interestingly and similar to the training (GSE22138) dataset, 82% of mHYP2 (monosomy-enriched) group in the validation cohort underwent metastasis during follow-up, compared to only 11% of the mHYP5 (disomy/partial monosomy-enriched) group patients. In addition, 33% of intermediate prognostic mHYP4 (undetermined) and 44% mixed prognostic mHYP1 (immune-enriched) patients experienced metastasis. With increased frequency of mHYP3 (neural-enriched) community, we observed that it has poor overall survival and 57% of the mHYP3 samples were undergoing metastasis (Figure 6B). Overall, this identifies and validates novel uveal melanoma subtype communities and their prognostic significance.

##### Comparison of subtype communities to known uveal melanoma classes

Previously, transcriptomic subtypes of uveal melanoma have been defined by clustering of gene expression profiles. Two classes were discovered – class 1, with good prognosis and association with chromosome 3 disomy; and class 2, with poor prognosis, associated with chromosome 3 monosomy and metastasis [25-27]. To reconcile these communities with the gene expression subtypes, we checked for gene set enrichment of the gene signatures [26] for class 1 and class 2 uveal melanomas in this cohort. The class 2 signature was enriched and borderline enriched in the mHYP1 community (immune-enriched; FDR < 0.001; Figure 6C) and mHYP2 (monosomy; FDR = 0.27; Figure 6D) groups, respectively, whereas, unexpectedly, the class 1 signature was not significantly enriched in any other group. This may indicate that the class 1 signature may be a heterogeneous set of patients who are not confined to any of our given community. Overall, this suggest that our novel uveal melanoma subtype communities reveal additional heterogeneity with clinical significance that requires further investigation.

## Conclusions

These results demonstrate that no one clustering algorithm should be relied on to produce clusters which are robust and capture all heterogeneity in a dataset. Instead, multiple algorithms should be applied to the same dataset, and their results compared and reconciled. Our polyCluster tool provides a straightforward interface to cluster datasets using multiple algorithms, provides statistics on the quality of each clustering, and allows the user to fully understand how each result is related through multiple reconciliations. The demonstration that some low-frequency clusters – which may be lost or discarded as outliers if only one algorithm is applied – are consistently identified across algorithms lends credence to their validity, and here such communities were additionally validated in an independent dataset. Thus, the reconciliation of multiple clustering results enables finer stratification of patients’ molecular profiles enabling more focused biological profiling.

## Methods

### Datasets

The breast cancer dataset [22] consists of 118 gene expression profiles generated from frozen resected samples. Patients in this were mostly early-stage, and were a mixture of node- and ER-positive and -negative. The discovery uveal melanoma dataset (GSE22138 [28]) consists of gene expression profiles for 63 untreated patients, chromosome 3 monosomy status and follow-up metastasis-free survival information. The validation dataset (GSE44295 [30]) contains 58 gene expression profiles from enucleation specimens, with metastasis-free survival information.

### Finding the optimal number of clusters

It is a not optimal for each of the above clustering methods to find local solutions which depend on the initial conditions, rather than robust clusterings that are stable over various input parameters. To address this, consensus clustering approaches repeat several iterations of the same algorithm using different random starting points, and can also perform the clustering over different subsets of samples. Consensus clustering for each algorithm was performed over a range of *k*-values from 2 to 10 and over multiple subsets of the data. The results of the consensus clustering were then inspected in order to determine the optimal *k*. Determining the optimal *k* from visual inspection alone is subjective, and so quantification of the consensus clustering is required. Here, the cophenetic correlation coefficient [31] and the silhouette width [21] were used to score each clustering.

### Hypergeometric test

Previous works have used the hypergeometric test to determine if different algorithms’ subtypes correspond to one another [10]. In this pipeline, comparisons can be made between any number of clustering algorithms. The hypergeometric test based false discovery rate (FDR) indicating the significance of the size of the overlap between two clusters was used.

#### Statistical analysis

FDR values for enrichment of gene sets were reported as calculated by the Broad Institute’s GSEA software [32]. Kaplan-Meier analysis was used to assess survival and p-values determined from the log-rank test. PAM analysis to generate centroids and assign subtypes using Pearson correlation and gene expression data was done as previously described [11].

#### Software

Code for hierarchical and k-means consensus clustering was adapted from the *ConsensusClusterPlus* v1.36.0 [33] R package. NMF was performed via the *nmf*v0.20.6 R package [34]. The *igraph* R package v1.0.1[35] was used for plotting networks and community detection. *Silhouette* width was calculated and plotted using the *silhouette* function from the R package *cluster* v2.0.4 [36]. Survival analysis was performed using the *survival* v2.39-5 R package [37]. GSEA was performed using the Broad Institute GSEA software [32]. The pipeline described in this paper is publicly available on GitHub at https://github.com/syspremed/polyClustR.

## Declarations

### Availability of data and material

Gene expression data analysed in this study are publicly available from the original publications (breast cancer data [22] and uveal melanoma [28], [30]) and through ArrayExpress with access number E-TABM-158 (breast) and Gene Expression Omnibus (GEO) with accession numbers GSE22138 and GSE44295 (uveal melanoma).

### Authors’ contributions

KE wrote the manuscript, developed the polyCluster package, performed all the experiments and analysed the results. GN helped with the statistical methods and oversaw the data analysis. AS conceived the idea, interpreted the results and wrote the manuscript.

## Acknowledgments

We acknowledge NHS funding to the NIHR Biomedical Research Centre at The Royal Marsden and the ICR.

